# Antivirals against monkeypox infections

**DOI:** 10.1101/2023.04.19.537483

**Authors:** Kevin Chiem, Aitor Nogales, Maria Lorenzo, Desarey Morales Vasquez, Yan Xiang, Yogesh K. Gupta, Rafael Blasco, Juan Carlos de la Torre, Luis Martínez-Sobrido

## Abstract

Monkeypox virus (MPXV) infection in humans are historically restricted to endemic regions in Africa. However, in 2022, an alarming number of MPXV cases have been reported globally with evidence of person-to-person transmission. Because of this, the World Health Organization (WHO) declared the MPXV outbreak a public health emergency of international concern. MPXV vaccines are limited and only two antivirals, tecovirimat and brincidofovir, approved by the United States (US) Food and Drug Administration (FDA) for the treatment of smallpox, are currently available for the treatment of MPXV infection. Here, we evaluated 19 compounds previously shown to inhibit different RNA viruses for their ability to inhibit Orthopoxvirus infections. We first used recombinant vaccinia virus (rVACV) expressing fluorescence (Scarlet or GFP) and luciferase (Nluc) reporter genes to identify compounds with anti-Orthopoxvirus activity. Seven compounds from the ReFRAME library (antimycin A, mycophenolic acid, AVN- 944, pyrazofurin, mycophenolate mofetil, azaribine, and brequinar) and six compounds from the NPC library (buparvaquone, valinomycin, narasin, monensin, rotenone, and mubritinib) showed antiviral activity against rVACV. Notably, the anti-VACV activity of some of the compounds in the ReFRAME library (antimycin A, mycophenolic acid, AVN- 944, mycophenolate mofetil, and brequinar) and all the compounds from the NPC library (buparvaquone, valinomycin, narasin, monensin, rotenone, and mubritinib) were confirmed with MPXV, demonstrating the broad-spectrum antiviral activity against Orthopoxviruses and their potential to be used for the antiviral treatment of MPXV, or other Orthopoxvirus, infections.

**IMPORTANCE:** Despite the eradication of smallpox, some Orthopoxviruses remain important human pathogens, as exemplified by the recent 2022 monkeypox virus (MPXV) outbreak. Although smallpox vaccines are effective against MPXV, there is presently limited access to those vaccines. In addition, current antiviral treatment against MPXV infections is limited to the use of the FDA-approved drugs tecovirimat and brincidofovir. Thus, there is an urgent need to identify novel antivirals for the treatment of MPXV, and other potentially zoonotic Orthopoxvirus infections. Here, we show that thirteen compounds, derived from two different libraries, previously found to inhibit several RNA viruses, exhibit also antiviral activity against VACV. Notably, eleven compounds also displayed antiviral activity against MPXV, demonstrating their potential to be incorporated into the therapeutic armamentarium to combat Orthopoxvirus infections.

## INTRODUCTION

In 2022, the World Health Organization (WHO) declared the monkeypox virus (MPXV) outbreak a public health emergency of international concern (1-3). Historically, MPXV infections in humans are rarely reported outside its endemic regions in Africa (4), but recently, in 2022, MPXV cases have been reported worldwide with sustained person-to-person transmission (1). Most of the confirmed cases in nonendemic regions are in Europe and North America (2), with no clear epidemiological links to endemic countries (5-7). As of March 2023, over 30,286 confirmed cases and 38 deaths related to MPXV infections have been reported in the United States (US) (8).

MPXV belongs to the Orthopoxvirus genus of the poxvirus family which includes variola virus (VARV), cowpox, and vaccinia virus (VACV) (9, 10). Poxviruses are large double stranded DNA viruses with a genome ranging from 135 to 380 kb encoding for up to 328 predicted open reading frames (ORF) (11). Currently, three clades of MPXV have been recognized: clade I predominant in the Congo Basin and responsible of up to 10% lethality in humans, clade IIa in West Africa responsible of low mortality in humans, and clade IIb, responsible of the currently global spreading in humans (12). To date, all MPXV cases associated with the 2022 global outbreak appeared to be related to clade II (13, 14). MPXV was first discovered in research monkeys in an animal facility in Copenhagen, Denmark, that were shipped from Singapore in 1958 (15, 16). Despite its name, the natural reservoir of MPXV clades I and IIa is not nonhuman primates, and instead it has been speculated to be small rodents native to Central and West Africa, respectively (17-21). Zoonotic transmission of MPXV most likely is mediated by body fluids, such as salivary or respiratory droplets, or derived from wounds (9, 22, 23. Person-to-person transmission of MPXV occurs by prolonged close contact exposure, for instance, prolonged face-to-face or intimate physical contact, touching infectious lesions or bodily fluids, and contaminated fomites (23-25).

Currently, two vaccines are available for the prophylactic treatment of MPXV infection, a modified vaccinia Ankara (MVA; JYNNEOS in the US, IMVANEX in the European Union, and IMAMUNE in Canada) (26-33) and ACAM2000 (34-36); the latter available for use under an Expanded Access Investigational New Drug (EA-IND) protocol. However, vaccine supplies are limited and are reserved for individuals that are of high risk, including immunocompromised or men that are sexually active with men (37). The only available FDA-approved treatments against poxviruses are tecovirimat and brincidofovir (38-41). Tecovirimat has been shown to target the VP37 protein (the product of F13L gene) of VACV that is required for extracellular virus particle formation and is highly conserved in the Orthopoxvirus genus (42, 43). However, resistance mutations to tecovirimat have been reported for various Orthopoxviruses, including VACV, camelpox, and cowpox (38, 44). Due to the severity of the MPXV outbreak, brincidofovir is available under the EA-IND (45), but its efficacy has not been fully defined outside animal models (46-53). Therefore, the importance of developing new therapeutics targeting host factors or virus-host interactions to combat Orthopoxvirus infections treatment can not be underestimated. The discovery and implementation of new antivirals is labor intense and costly process requiring multiple rounds of testing and refinement to ensure their safety and efficacy. Repurposing current US FDA-approved drugs can facilitate the advancement of candidate antiviral drugs into the clinic, since the pharmacology and toxicology of the drug have already been established.

In this study, we used a previously described bireporter cell-based assay based on the use of recombinant vaccinia virus (rVACV) expressing fluorescent (GFP or Scarlet) and luciferase (Nluc) proteins (54) to assess the antiviral efficacy of compounds previously identified as having antiviral activity against different RNA viruses. These compounds belonged to two libraries: Repurposing, Focused Rescue, and Accelerated Medchem (ReFRAME) (55-58) and NCATS Pharmaceutical Collection (NPC) (59). We found that seven of the compounds from the ReFRAME library (antimycin A, mycophenolic acid, AVN-944, pyrazofurin, mycophenolate mofetil, azaribine, and brequinar) and six compounds from the NPC library (buparvaquone, valinomycin, narasin, monensin, rotenone, and mubritinib) had antiviral activity against VACV. Importantly the broad anti-Orthopoxvirus activity of some of the identified compounds was confirmed with MPXV. These results open the prospect of implementing these, or related, compounds for the treatment of Orthopoxvirus infection, including MPXV, which would add new compounds to the therapeutic armamentarium currently available to combat Orthopoxvirus infections.

## MATERIALS AND METHODS

### Biosafety

Experiments with MPXV were performed at biosafety level 3 (BSL3) containment laboratories at Texas Biomedical Research Institute (TX Biomed) and were approved by the Institutional Biosafety Committee (IBC) at TX Biomed. All the researchers involved in studies using MPXV were vaccinated with the modified vaccinia Ankara JYNNEOS vaccine.

### Cell lines

African green monkey kidney fibroblast (CV-1; ATCC CCL-70) and human adenocarcinoma alveolar basal epithelial (A549; ATCC CCL-185) cells were maintained in Dulbecco’s modified Eagle’s medium (DMEM; Corning) containing 5% fetal bovine serum (FBS) and 1% PSG (100 U/ml penicillin, 100 µg/ml streptomycin and 2 mM L-glutamine). Cells were grown at 37°C in a 5% CO_2_ atmosphere.

### Viruses

The two reporter-expressing recombinant (r)VACV expressing fluorescent (GFP or Scarlett) and luciferase (Nluc) proteins, rVACV Nluc/GFP and rVACV Nluc/Scarlet, respectively were previously described and characterized (54). We used both fluorescent-expressing rVACV to assess the feasibility of using both GFP and Scarlet coupled with fluorescent microscopy and plate readers to identify compounds with antiviral activity (54). We selected Nluc since is a secreted small luciferase that allow us to detect bioluminescence activity in cell culture supernatants and, therefore, used in longitudinal studies (54). MPXV USA-2003 (NR-2500) clade II was obtained from the Biodefense and Emerging Infectious (BEI) Resources repository and propagated in CV- 1 cells. Virus infections were conducted in DMEM containing 2% FBS and 1% PSG.

### Compounds

The compounds previously identified to have antiviral activity against RNA viruses in the ReFRAME library were previously described (55-58) and includes antimycin A (Sigma-Aldrich, Cat. # A8674), OSU-03012 (AkSci, Cat. # Y0267), mycophenolic acid (AkSci, Cat. # E480), AVN-944 (ADOOQ Bio, Cat. # A13652), 6-azauridine (azauridine; Sigma-Aldrich, Cat. # A1882), pyrazofurin (Sigma-Aldrich, Cat. # SLM1502), mycophenolate mofetil (AkSci, Cat. # J90063), 2’,3’,5’-Triacetyl-6-azauridine (azaribine; Sigma-Aldrich, Cat. # T340057), and brequinar sodium salt hydrate (brequinar; Sigma-Aldrich, Cat. # SML0113). Likewise, the compounds in the NPC library previously shown to have antiviral activity against RNA viruses were previously described (59) and include azoxystrobin (Sigma-Aldrich, Cat. # 31697), buparvaquone (ArkPharm, Cat. # 88426-33-9), valinomycin (Sigma-Aldrich, Cat. # V0627), narasin sodium salt (narasin; Cayman Chemical, Cat. # 19447), amuvatinib (MedChemExpress, Cat. # HY-10206), monensin sodium salt (monensin; Sigma-Aldrich, Cat. # M5273), spautin-1 (MedChemExpress, Cat. # HY-12990), tryptanthrin (Sigma-Aldrich, Cat. # SML0310), rotenone (Sigma-Aldrich, Cat. # 45656), and mubritinib (MedChemExpress, Cat. # HY- 13501). Tecovirimat and 2,2’-Bi-1H-benzimidazole (Benzimidazole; NSC-67061) were included as positive and negative controls, respectively in all the assays (41, 60). All compounds were prepared at 10 mM stock solution in dimethyl sulfoxide (DMSO) and kept at -20°C until experimentation. All compounds were diluted in DMEM supplemented with 2% FBS and 1% PSG (infection medium). The maximum concentration of DMSO in all antiviral preparations was 0.1%, including vehicle control wells.

### Bireporter cell-based assay for the identification of antivirals

Confluent monolayers of human A549 cells (96-well plate format, 4 x 10^4^ cells/well, and quadruplicates) were infected with 200 plaque forming units (PFU)/well of rVACV Nluc/Scarlet or rVACV Nluc/GFP for 1 h at 37°C. After virus absorption, the virus inoculum was removed, and cells were incubated with infection medium containing 3- fold serial dilutions of the indicated compounds (starting concentration of 50 µM for all compounds except for azoxystrobin, buparvaquone, tryptanthrin, and OSU-03012 [starting at 450 µM]; mycophenolic acid, mycophenolate mofetil, and azaribine [starting at 150 µM; and antimycin A, narasin, monensin, valinomycin and AVN-944 [starting at 1.85 µM]). Mock-infected cells and cells infected in the absence of drug were included as an internal control. At 24 h post-infection (hpi), fluorescence expression (GFP and Scarlet) was visualized using a fluorescence microscope and quantified using a microplate reader (BioTek Synergy). Simultaneously, cell culture supernatants from the same infections were collected and used to measure Nluc expression using the Nano-Glo luciferase substrate (Promega) and the microplate reader (BioTek Synergy). Percent of viral infection was determined based on fluorescent (GFP or Scarlet) and Nluc signals, as previously described (58, 61, 62. Experiments were conducted thrice using technical quadruplicates. Microsoft Excel was used to calculate the mean and standard deviation (SD) of viral inhibition from quadruplicate wells. Half maximal effective concentration (EC_50_) values were determined using sigmoidal dose response curves on GraphPad Prism (Version 9).

### Focus forming reduction assay (FFRA)

Confluent monolayers of human A549 cells (96-well plate format, 4 x 10^4^ cells/well, and quadruplicates) were infected with 200 PFU/well of MPXV for 1 h at 37°C. After virus absorption, the virus inoculum was removed, and cells were incubated with infection medium containing 3-fold serial dilutions (starting concentration of 50 µM for all compounds except for azoxystrobin, buparvaquone, tryptanthrin, and OSU-03012 [starting at 450 µM]; mycophenolic acid, mycophenolate mofetil, and azaribine [starting at 150 µM; and antimycin A, narasin, monensin, valinomycin and AVN-944 [starting at 1.85 µM]) of the indicated compounds and 1% Avicel (Sigma-Aldrich). Mock-infected cells and cells infected in the absence of drug were included as internal controls. At 24 hpi, cells were fixed in 10% neutral buffered formalin for 24 h, and then permeabilized with 0.5% Triton X-100 in PBS for 10 mins at room temperature (RT). Then, cells were blocked with 2.5% bovine serum albumin (BSA) in PBS for 1 h, followed by immunostaining with an anti-VACV A33R polyclonal antibody (BEI Resources, NR-628), Vectastain ABC kit, and DAB peroxidase substrate kit (Vector Laboratories), based on the manufacturer’s recommendations. Viral infections were determined based on the number of plaques present in each of the 96 well plates using an ImmunoSpot plate reader, as previously described (57, 63-65). Similar assays have been described in the literature to assess the antiviral activity of compounds (66, 67). Experiments were conducted thrice using technical quadruplicates. Microsoft Excel was used to calculate the mean and SD of viral inhibition from quadruplicate wells. Non-linear regression curves and EC_50_ values were determined using sigmoidal dose response curves on GraphPad Prism (Version 9).

### Viral titer reduction assays

Confluent monolayers of human A549 cells (24-well plate format, 2 x 10^5^ cells/well, triplicates) were infected with rVACV Nluc/Scarlet, rVACV Nluc/GFP, or MPXV at MOI of 0.01 or MOI of 3 for 1 h at 37°C. After virus absorption, the virus inoculum was removed, and cells were incubated with infection medium containing 10-fold serial dilutions of tecovirimat (starting concentration of 100 µM). Mock-infected cells and cells infected in the absence of drug were included as internal controls. At 24, 48, and 72 hpi, cell culture supernatant was collected, and Nluc activity was determined by adding Nano-Glo luciferase substrate (Promega) and quantified in a microplate reader (rVACV Nluc/Scarlet or rVACV Nluc/GFP). Viral titers in the cell culture supernatants were determined by plaque assay in CV-1 cells. Briefly, cells were infected with 10-fold serially diluted supernatants for 1 h. After viral adsorption, cells were overlayed with medium containing 1% Avicel (Sigma-Aldrich). Cells were fixed in 10% neutral buffered formalin for 24 h and then permeabilized with 0.5% Triton X-100 in PBS for 10 min at room temperature. Next, cells were blocked with 2.5% bovine serum albumin (BSA) in PBS for 1 h and immunostained with an anti-VACV A33R polyclonal antibody (BEI Resources, NR-628) and developed with an anti-rabbit Vectastain ABC kit, and DAB peroxidase substrate kit (Vector Laboratories), following manufacturer’s recommendations.

### Cell viability assays

MTT (CellTiter 96 nonradioactive cell proliferation assay, Promega) and XTT (cell viability and proliferation assay; Sigma-Aldrich) assays were used to determine A549 cell viability as previously described (58). Briefly, confluent monolayers of human A549 cells (96-well plate format, 4 x 10^4^ cells/well, quadruplicates) were incubated with 100 µl of infection medium containing 3-fold dilutions of compounds (starting concentration of 50 µM for all compounds except for azoxystrobin, buparvaquone, tryptanthrin, and OSU-03012 [starting at 450 µM]; mycophenolic acid, mycophenolate mofetil, and azaribine [starting at 150 µM; and antimycin A, narasin, monensin, valinomycin and AVN-944 [starting at 1.85 µM]) or with 0.1% DMSO vehicle control. Then, plates were incubated at 37°C with 5% CO_2_ for 48 h, and subsequently treated with 15 µl of dye solution for the MTT assay, or 50 µl of XTT labeling reagent for the XTT assay, and incubated at 37°C for an additional 4 h. Stop solution was then added to the MTT assay to halt the reaction. Absorbance (570 nm) in each of the wells was measured using a microplate reader (BioTek Synergy). Cell viability was determined as the relative percent to values of DMSO vehicle-treated cells. Non-linear regression curves and the median cytotoxic concentration (CC_50_) at 48 h were determined using GraphPad Prism software (Version 9).

### Statistical analysis

GraphPad Prism software (Version 9) was used for data analysis. CC_50_ and EC_50_ values were calculated using sigmoidal dose-response curves, and the selective index (SI) of each compound was determined by dividing the CC_50_ with the EC_50_ values. Significance was determined by standard unpaired Student’s *t*-test. Data of three independent experiments in quadruplicates were expressed as the mean and SD using Microsoft Excel software.

## RESULTS

### A bireporter cell-based assay for the identification of antivirals against VACV

To determine the antiviral activity of compounds from the ReFRAME (55-57) or NPC (59) libraries against Orthopoxviruses, which were identified as having antiviral activity against different RNA viruses, we used our previously described fluorescent (Scarlet or GFP) and Nluc expressing rVACV (rVACV Nluc/Scarlet and rVACV Nluc/GFP) (**Figure 1A**) (54). We have shown that expression levels of reporter genes from these rVACV can be used as an accurate surrogate of viral infection (54). First, we validated the use of reporter-expressing rVACV (rVACV Nluc/Scarlet and rVACV Nluc/GFP) by assessing the antiviral activity of tecovirimat, an FDA-approved antiviral against poxviruses (38-41). We used 3-fold serial dilutions (starting at 50 µM) of tecovirimat to treat human A549 cells infected with rVACV Nluc/Scarlet or rVACV Nluc/GFP and viral infection was assessed based on fluorescent and luciferase expression (**Figures 1B and C**). Tecovirimat exhibited a dose-dependent inhibitory effect in Scarlet (left) or GFP (right) expression levels (**Figure 1B**). Tecovirimat EC_50_ values for rVACV Nluc/Scarlet and Nluc/GFP were similar, 0.031 and 0.026 µM, respectively, regardless of which fluorescence reporter was used for quantifications (**Figure 1C, left panel**). Likewise, EC_50_ values calculated by assessing Nluc activity in the cell culture supernatants from rVACV Nluc/Scarlet or Nluc/GFP infected cells were also similar (0.047 and 0.033 µM, respectively) (**Figure 1C, right panel**). Selectivity index (SI) values based on Scarlet or GFP levels were also similar (>1,612.9 and >1,923.1, respectively) and comparable to those based on Nluc from either the rVACV Nluc/Scarlet or Nluc/GFP (>1,063.8 and >1,515.2, respectively). To further validate the use of bireporter rVACV to assess compound antiviral activity, we conducted a viral titer reduction assay. For this we infected A549 cells at low (0.01) and high (3) MOI and treated them with 10-fold serial dilutions of tecovirimat (starting concentration of 100 µM). Cell culture supernatants were collected at 24, 48, and 72 hpi. Then, extracellular viral titers and Nluc activity (rVACV Nluc/Scarlet or rVACV Nluc/GFP) were determined from the cell culture supernatants. We found a dose-dependent decrease in extracellular viral titers of rVACV Nluc/Scarlet, rVACV Nluc/GFP, and MPXV (**Figure 1D**). Tecovirimat primarily acts by inhibiting the activity of VACV protein VP37, which is required for virus egress. This results in decreased production of extracellular virus and viral spread without affecting gene expression, reflected in cell culture supernatants having Nluc activity even in the presence of the highest concentration (100 µM) of tecovirimat (**Figure 1E**). The decrease in extracellular viral titers in high MOI infections in the presence of Tecovirimat presumably reflects inhibition of extracellular virus formation by the drug. Nonetheless, the antiviral activity of tecovirimat obtained with rVACV Nluc/Scarlet and Nluc/GFP, or MPXV, was consistent with those reported in the literature for VACV (68) and MPXV (38, 39, 43, 69, 70). These results demonstrated that rVACV Nluc/Scarlet or rVACV Nluc/GFP can be used to determine accurately the inhibitory properties of antivirals against Orthopoxvirus based on fluorescent or luciferase expression.

**Figure 1.**
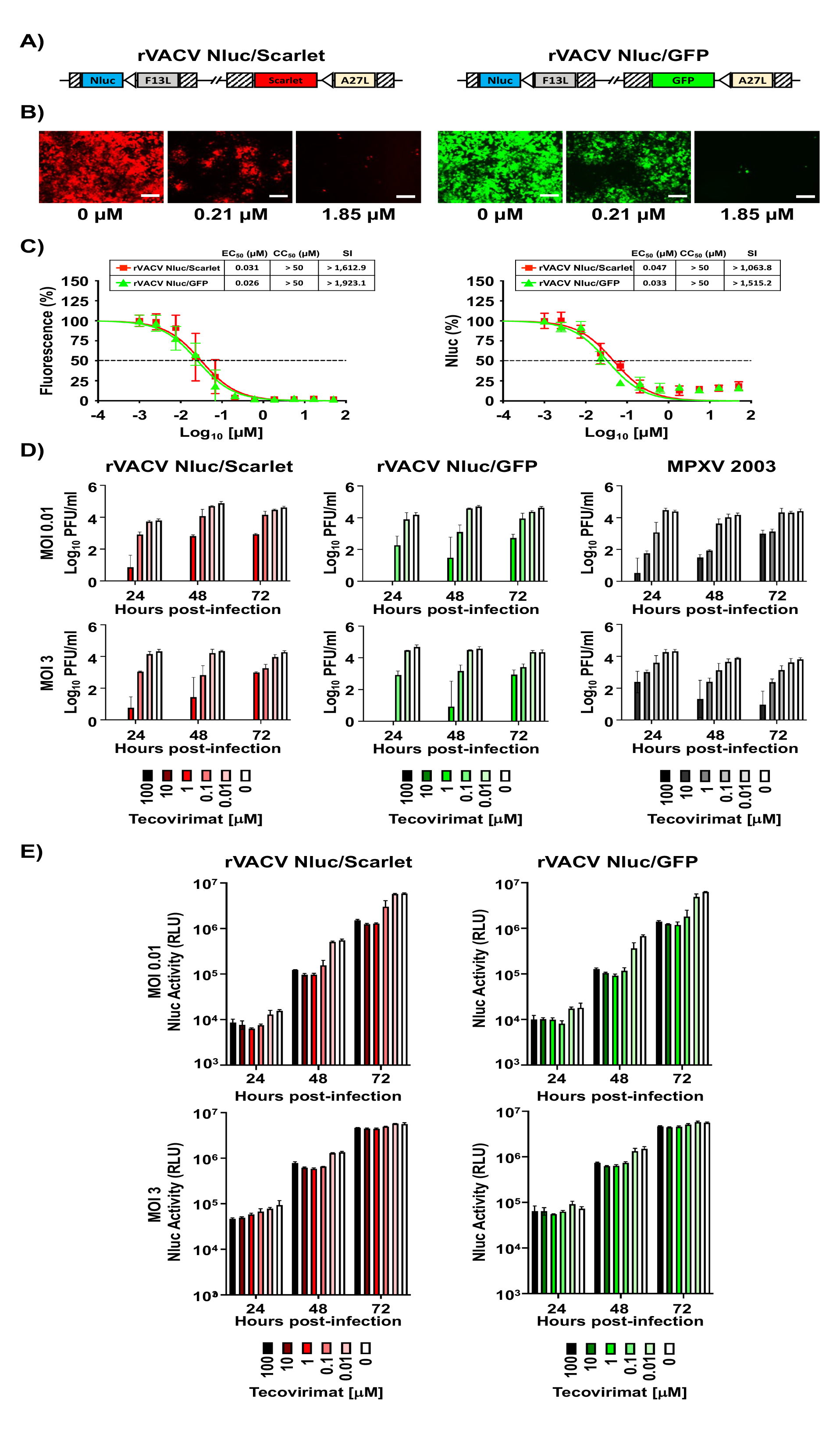
A bireporter cell-based assay for the identification of VACV antivirals. A) Schematic representation of rVACV Nluc/Scarlet (left) and rVACV Nluc/GFP (right): Nanoluciferase (Nluc, blue box) and fluorescent reporter Scarlet (Red box) or green fluorescent protein (GFP, green box) genes were inserted downstream of the F13L (gray box) or the A27L (yellow box), respectively, in the viral genome. The reporter genes are expressed by an early/late VACV synthetic promoter (white arrows). Striped boxes indicate the recombination flanking regions for the insertion of reporter genes into the VACV genome. **B-C) A bireporter cell-based assay to identify antivirals:** Human A549 cells (2×10^4^ cells/well, 96-well plates, quadruplicates) were infected with 200 PFU of rVACV Nluc/Scarlet (left) or rVACV Nluc/GFP (right) and incubated with 3-fold serial dilutions (starting concentration of 50 µM) of tecovirimat. Mock-infected cells and cells infected in the absence of drug were included as an internal control. At 24 hpi, Scarlet (rVACV Nluc/Scarlet, left) or GFP (rVACV Nluc/GFP, right) expression were visualized under a fluorescence microscope **(B).** Representative images are shown. Scale bars, 100 µm. Magnification, x20. To quantify inhibition of viral replication, Scarlet (red squares) or GFP (green triangles) expression levels were quantified at 24 hpi using a fluorescent microplate reader **(C, left panel)**; or by Nluc expression **(C, right panel)** using a luminometer for rVACV Nluc/Scarlet (red squares) or rVACV Nluc/GFP (green triangles). The EC_50_ of tecovirimat was calculated using sigmoidal dose-response curves. The CC_50_ of Tecovirimat was determined using a MTT assay kit. The SI was calculated by dividing the CC_50_/EC_50_. The percent of viral inhibition was normalized to non-tecovirimat-treated controls. The dotted lines indicate 50% inhibition. Data represent the means and standard deviations (SD) of the experiment conducted in quadruplicates. **D-E) Viral titers (D) and Nluc activity (E):** Human A549 cells (2×10^4^ cells/well, 96-well plates, quadruplicates) were infected with an MOI of 0.01 (top) or 3 (bottom) of rVACV Nluc/Scarlet, rVACV Nluc/GFP, or MPXV for 1 h, and incubated with 10-fold serial dilutions (starting concentration of 100 µM) of tecovirimat. At 24, 48, and 72 hpi, tissue culture supernatants were collected, and viral titers were determined by plaque assay in CV1 cells. Red bars (left), rVACV Nluc/Scarlet; green bars (center), rVACV Nluc/GFP; gray bars (right), MPXV. Additionally, Nluc activity (E) in cell culture supernatants were determined using a luminometer for rVACV Nluc/Scarlet (red bars, left) or rVACV Nluc/GFP (green bars, right). Mock-infected cells and cells infected in the absence of drug were included as internal controls.

### Effect of selected compounds from the ReFRAME and NPC libraries on VACV multiplication

To assess whether ReFRAME and NPC compounds previously identified to have antiviral activity against arenaviruses, influenza A and B viruses (IAV and IBV, respectively), and Zika virus (ZIKV) (55-57, 59, 64) also inhibited poxvirus infection, we use the above described bireporter cell-based assay based on the use of rVACV Nluc/Scarlet and rVACV Nluc/GFP. We included benzimidazole and tecovirimat as negative and positive controls, respectively. Seven of the compounds tested from the ReFRAME library showed potent inhibition of rVACV Nluc/Scarlet and Nluc/GFP based on fluorescence (**Figure 2A-I, top panels**) and Nluc (**Figure 2A-I, bottom panels**) activities (**Table 1**). As expected, we did not observe any antiviral activity in cells treated with benzimidazole (**Figure 2J and Table 1**), while tecovirimat antiviral activity (**Figure 2K and Table 1**) was consistent with the results of our validation experiment (**Figure 1**) and those reported in the literature (38, 39, 43, 69, 70). Antimycin A showed potent antiviral activity based on fluorescence (EC_50_ Scarlet = 0.07 µM; EC_50_ GFP = 0.06 µM) or Nluc (EC_50_ Scarlet = 0.09 µM; EC_50_ GFP = 0.12 µM) expression. However, the SI value of antimycin A differed based on the CC_50_ results from the MTT or XTT toxicity assays (**Table 1**). AVN-944 also exhibited potent inhibitory properties against VACV as determined by fluorescence (EC_50_ Scarlet = 0.13 µM; EC_50_ GFP = 0.07 µM) or Nluc (EC_50_ Scarlet = 0.11 µM; EC_50_ GFP = 0.07 µM) expression. In the case of AVN-944, the SI value was not altered by the CC_50_ results from the MTT or XTT toxicity assays (**Table 1**). Other compounds from the ReFRAME library also showed inhibition of rVACV, including mycophenolic acid, pyrazofurin, mycophenolate mofetil, azaribine, and brequinar (**Figure 2 and Table 1**). Likewise, six compounds from the NPC library (buparvaquone, valinomycin, narasin, monensin, rotenone, and mubritinib), previously found to have anti-arenaviral activity (59), showed potent inhibition of rVACV Nluc/Scarlet and Nluc/GFP based on fluorescence (**Figure 3A-J, top panels**) and Nluc (**Figure 3A-J, bottom panels**) expression (**Table 2**). As above, benzimidazole (**Figure 3J and Table 2**) and tecovirimat (**Figure 3K and Table 2**) treatment had none and potent, respectively, antiviral activity. The most potent antiviral compounds against VACV in the NPC library were valinomycin, rotenone, and mubritinib, followed by monensin, narasin, and buparvaquone (**Table 2**). In the case of the compounds from the NPC library, the SI values, except monensin, were different based on the CC_50_ results from the MTT or XTT toxicity assays (**Table 2**). These results demonstrated that compounds in the ReFRAME (**Figure 2 and Table 1**) and NPC (**Figure 3 and Table 2**) libraries previously identified as having antiviral activity against different RNA viruses (55-57, 59) also have potent antiviral activity against VACV, demonstrating their broad-spectrum antiviral activity against both RNA and DNA viruses.

**Figure 2.**
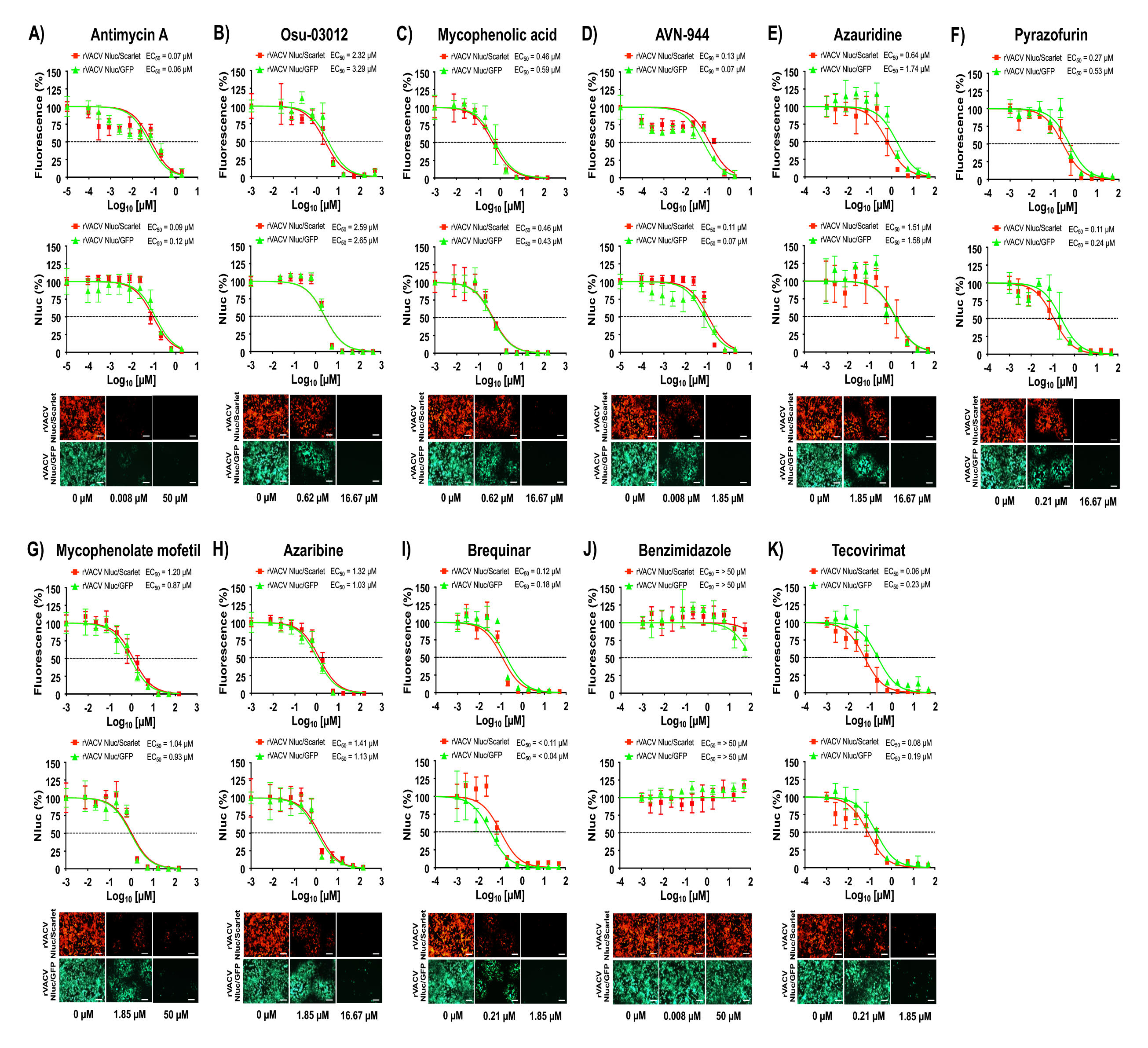
Antiviral activity of the ReFRAME antiviral compounds against VACV: Human A549 cells (2×10^4^ cells/well, 96-well plates, quadruplicates) were infected with 200 PFU of rVACV Nluc/Scarlet (red squares) or rVACV Nluc/GFP (green triangles) and incubated with 3-fold serial dilutions of antimycin A (**A**), Osu-03012 (**B**), mycophenolic acid (**C**), AVN-944 (**D**), azauridine (**E**), pyrazofurin (**F**), mycophenolate mofetil (**G**), azaribine (**H**), or brequinar (**I**). Benzimidazole (**J**) and tecovirimat (**K**) were included as negative and positive controls, respectively. Mock-infected cells and cells infected in the absence of drug were included as an internal control. At 24 hpi, inhibition of rVACV Nluc/Scarlet and rVACV Nluc/GFP viral replication were evaluated by quantifying fluorescent Scarlet or GFP (top graphs), or Nluc (bottom graphs) expression using a fluorescent microplate reader and a luminometer, respectively. The EC_50_ for each compound was calculated using sigmoidal dose-response curves with GraphPad Prism. The dotted lines indicate 50% inhibition. Data represent the means and SD from quadruplicates. At the same hpi, Scarlet (rVACV Nluc/Scarlet, top) and GFP (rVACV Nluc/GFP, bottom) fluorescent expression in infected cells in the absence (0 µM) or in the presence of the indicated concentrations of the antiviral compounds (max concentration and EC_50_) were visualized using a fluorescence microscope. Representative images are shown. Scale bars, 100 µm. Magnification, x20.

**Figure 3.**
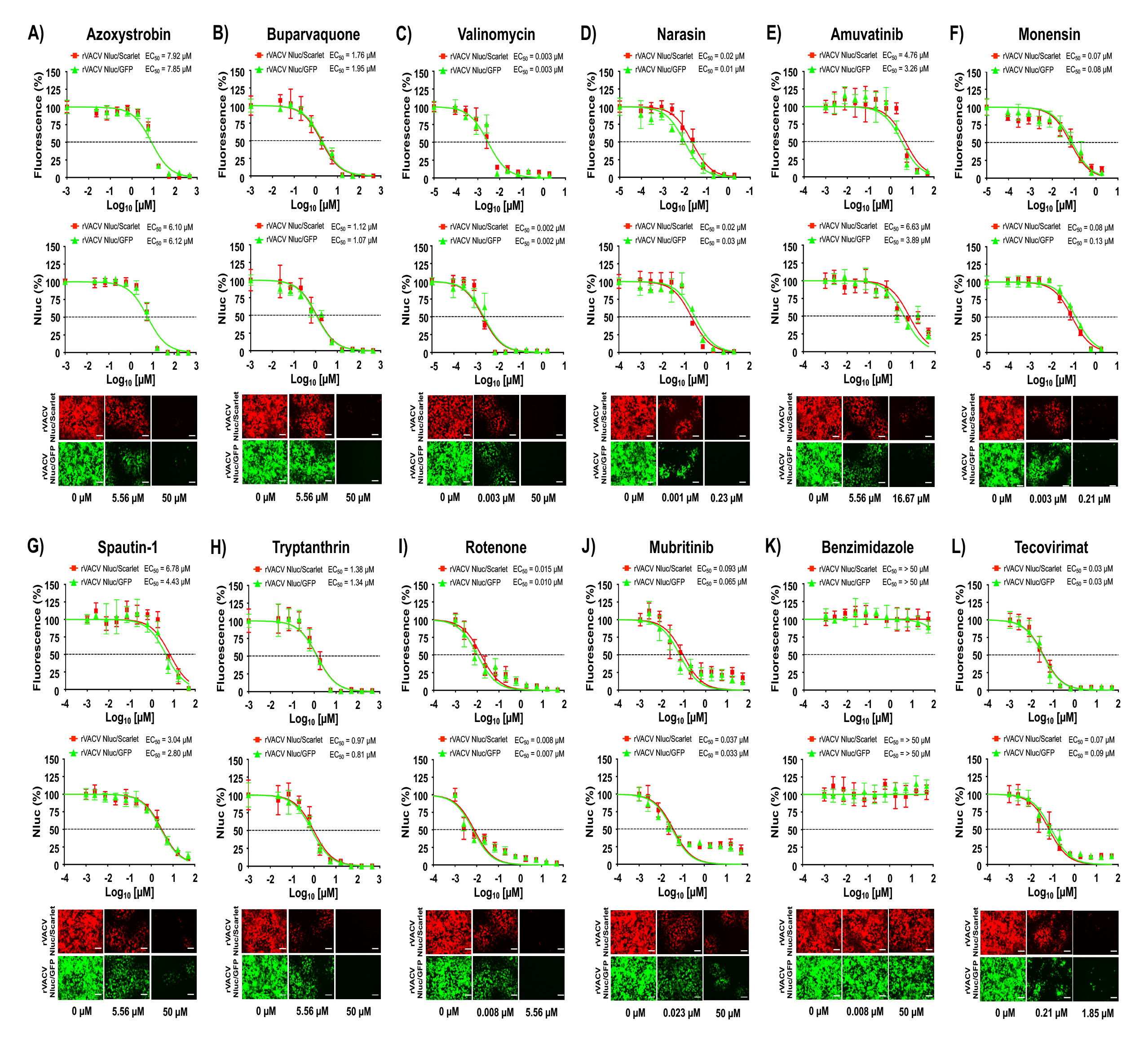
Antiviral activity of the NPC antiviral compounds against VACV: Human A549 cells (2×10^4^ cells/well, 96-well plates, quadruplicates) were infected with 200 PFU of rVACV Nluc/Scarlet (red squares) or rVACV Nluc/GFP (green triangles) and incubated with 3-fold serial dilutions of azoxystrobin (**A**), buparvaquone (**B**), valinomycin (**C**), narasin (**D**), amuvatinib (**E**), monensin (**F**), spautin-1 (**G**), tryptanthrin (**H**), rotenone (**I**), or mubritinib (**J**) Benzimidazole (**K**) and tecovirimat (**L**) were included as negative and positive controls, respectively. Mock-infected cells and cells infected in the absence of drug were included as an internal control. Inhibition of rVACV Nluc/Scarlet or rVACV Nluc/GFP viral replication were evaluated by quantifying Scarlet or GFP (top graphs) and Nluc (bottom graphs) expression at 24 hpi using fluorescent and luciferase microplate reader, respectively. The EC_50_ for each compound was calculated using sigmoidal dose-response curves with GraphPad Prism. The dotted lines indicate 50% inhibition. Data represent the means and SD from quadruplicates. At same hpi, Scarlet (rVACV Nluc/Scarlet; top) and GFP (rVACV Nluc/GFP; bottom) fluorescent expression in infected cells in the absence (0 µM) or in the presence of the indicated concentrations of the antiviral compounds (max concentration and EC_50_) were visualized using a fluorescence microscope. Representative images are shown. Scale bars, 100 µm. Magnification, x20.

**Table 1.** CC_50_, EC_50_, and SI of the ReFRAME antiviral compounds against the bireporter rVACV.

| Compound | Virus | CC<sub>50</sub> (μM) |  | EC<sub>50</sub> (μM) |  | SI (FP) |  | SI (Nluc) |  |
| --- | --- | --- | --- | --- | --- | --- | --- | --- | --- |
|  |  | MTT | XTT | FP | Nluc | MTT | XTT | MTT | XTT |
| Antimycin A | rVACV Scarlet/Nluc | 8.73 | 39.29 | 0.07 | 0.09 | 125 | 561 | 97 | 437 |
|  | rVACV GFP/Nluc | 8.73 | 39.29 | 0.06 | 0.12 | 146 | 655 | 73 | 327 |
| Osu-03012 | rVACV Scarlet/Nluc | 22.23 | 43.72 | 2.32 | 2.59 | 10 | 19 | 9 | 17 |
|  | rVACV GFP/Nluc | 22.23 | 43.72 | 3.29 | 2.65 | 7 | 13 | 8 | 16 |
| Mycophenolic acid | rVACV Scarlet/Nluc | > 150 | > 150 | 0.46 | 0.46 | > 326 | > 326 | > 326 | > 326 |
|  | rVACV GFP/Nluc | > 150 | > 150 | 0.59 | 0.43 | > 254 | > 254 | > 349 | > 349 |
| AVN-944 | rVACV Scarlet/Nluc | > 50 | > 50 | 0.13 | 0.11 | > 385 | > 385 | > 455 | > 455 |
|  | rVACV GFP/Nluc | > 50 | > 50 | 0.07 | 0.07 | > 714 | > 714 | > 714 | > 714 |
| Azauridine | rVACV Scarlet/Nluc | > 50 | > 50 | 0.64 | 1.51 | > 78 | > 78 | > 33 | > 33 |
|  | rVACV GFP/Nluc | > 50 | > 50 | 1.74 | 1.58 | > 29 | > 29 | > 32 | > 32 |
| Pyrazofurin | rVACV Scarlet/Nluc | > 50 | > 50 | 0.27 | 0.11 | > 185 | > 185 | > 455 | > 455 |
|  | rVACV GFP/Nluc | > 50 | > 50 | 0.53 | 0.24 | > 94 | > 94 | > 208 | > 208 |
| Mycophenolate mofetil | rVACV Scarlet/Nluc | > 150 | > 150 | 1.20 | 1.04 | > 125 | > 125 | > 144 | > 144 |
|  | rVACV GFP/Nluc | > 150 | > 150 | 0.87 | 0.93 | > 172 | > 172 | > 161 | > 161 |
| Azaribine | rVACV Scarlet/Nluc | > 150 | > 150 | 1.32 | 1.41 | > 114 | > 114 | > 106 | > 106 |
|  | rVACV GFP/Nluc | > 150 | > 150 | 1.03 | 1.13 | > 146 | > 146 | > 133 | > 133 |
| Brequinar | rVACV Scarlet/Nluc | > 50 | > 50 | 0.12 | 0.11 | > 417 | > 417 | > 455 | > 455 |
|  | rVACV GFP/Nluc | > 50 | > 50 | 0.18 | 0.04 | > 278 | > 278 | > 1,250 | > 1,250 |
| Benzimidazole | rVACV Scarlet/Nluc | > 50 | > 50 | > 50 | > 50 | > 1 | > 1 | > 1 | > 1 |
|  | rVACV GFP/Nluc | > 50 | > 50 | > 50 | > 50 | > 1 | > 1 | > 1 | > 1 |
| Tecovirimat | rVACV Scarlet/Nluc | > 50 | > 50 | 0.06 | 0.08 | > 833 | > 833 | > 625 | > 625 |
|  | rVACV GFP/Nluc | > 50 | > 50 | 0.23 | 0.19 | > 217 | > 217 | > 263 | > 263 |

FP = fluorescent protein

**Table 2.** CC_50_, EC_50_, and SI of the NPC antiviral compounds against the bireporter rVACV.

| Compound | Virus | CC<sub>50</sub> (μM) |  | EC<sub>50</sub> (μM) |  | SI (FP) |  | SI (Nluc) |  |
| --- | --- | --- | --- | --- | --- | --- | --- | --- | --- |
|  |  | MTT | XTT | FP | Nluc | MTT | XTT | MTT | XTT |
| Azoxystrobin | rVACV Scarlet/Nluc | 27.92 | > 450 | 7.92 | 6.10 | 4 | > 57 | 5 | > 74 |
|  | rVACV GFP/Nluc | 27.92 | > 450 | 7.85 | 6.12 | 4 | > 57 | 5 | > 74 |
| Buparvaquone | rVACV Scarlet/Nluc | 121.4 | > 450 | 1.76 | 1.12 | 69 | > 256 | 108 | > 402 |
|  | rVACV GFP/Nluc | 121.4 | > 450 | 1.95 | 1.07 | 62 | > 231 | 113 | > 421 |
| Valinomycin | rVACV Scarlet/Nluc | 0.29 | 13.63 | 0.003 | 0.002 | 97 | 4,543 | 145 | 6,815 |
|  | rVACV GFP/Nluc | 0.29 | 13.63 | 0.003 | 0.002 | 97 | 4,543 | 145 | 6,815 |
| Narasin | rVACV Scarlet/Nluc | 1.3 | 8.73 | 0.02 | 0.02 | 65 | 437 | 65 | 437 |
|  | rVACV GFP/Nluc | 1.3 | 8.73 | 0.01 | 0.03 | 130 | 873 | 43 | 291 |
| Amuvatinib | rVACV Scarlet/Nluc | 18.51 | 1.80 | 4.76 | 6.63 | 4 | 0 | 3 | 0 |
|  | rVACV GFP/Nluc | 18.51 | 1.80 | 3.26 | 3.89 | 6 | 1 | 5 | 0 |
| Monensin | rVACV Scarlet/Nluc | > 50 | > 50 | 0.07 | 0.08 | > 714 | > 714 | > 625 | > 625 |
|  | rVACV GFP/Nluc | > 50 | > 50 | 0.08 | 0.13 | > 625 | > 625 | > 385 | > 385 |
| Spautin-1 | rVACV Scarlet/Nluc | 8.00 | > 50 | 6.78 | 3.04 | 1 | > 7 | 3 | > 16 |
|  | rVACV GFP/Nluc | 8.00 | > 50 | 4.43 | 2.80 | 2 | > 11 | 3 | > 18 |
| Tryptanthrin | rVACV Scarlet/Nluc | 6.31 | 16.65 | 1.38 | 0.97 | 5 | 12 | 7 | 17 |
|  | rVACV GFP/Nluc | 6.31 | 16.65 | 1.34 | 0.81 | 5 | 12 | 8 | 21 |
| Rotenone | rVACV Scarlet/Nluc | 0.10 | > 50 | 0.015 | 0.008 | 7 | > 3,333 | 13 | > 6,250 |
|  | rVACV GFP/Nluc | 0.10 | > 50 | 0.010 | 0.007 | 10 | > 5,000 | 14 | > 7,143 |
| Mubritinib | rVACV Scarlet/Nluc | 0.07 | > 50 | 0.093 | 0.037 | 1 | > 538 | 2 | > 1,351 |
|  | rVACV GFP/Nluc | 0.07 | > 50 | 0.065 | 0.033 | 1 | > 769 | 2 | > 1,515 |
| Benzimidazole | rVACV Scarlet/Nluc | > 50 | > 50 | > 50 | > 50 | > 1 | > 1 | > 1 | > 1 |
|  | rVACV GFP/Nluc | > 50 | > 50 | > 50 | > 50 | > 1 | > 1 | > 1 | > 1 |
| Tecovirimat | rVACV Scarlet/Nluc | > 50 | > 50 | 0.03 | 0.07 | > 1,667 | > 1,667 | > 714 | > 714 |
|  | rVACV GFP/Nluc | > 50 | > 50 | 0.03 | 0.09 | > 1,667 | > 1,667 | > 556 | > 556 |

FP = fluorescent protein

### Effect of selected compounds from the ReFRAME and NPC libraries on MPXV multiplication

VACV and MPXV are both Orthopoxviruses and share similar genetic and biological properties. We therefore examined whether compounds with anti-VACV activity from the ReFRAME and NPC libraries were also able to inhibit MPXV infection. To assess the ability of the compounds in the ReFRAME and NPC libraries to inhibit MPXV we used a focus forming reduction assay (FFRA). We found that five of the compounds tested from the ReFRAME library (**Figure 4A and Table 3**) and all of the compounds from the NPC library (**Figure 4B and Table 4**) also had antiviral activity against MPXV. As anticipated, benzimidazole (**Figures 4A and 4B**) did not inhibit MPXV infection, whereas tecovirimat (**Figures 4A and 4B**) potently inhibited MPXV (71). As with VACV, antimycin A, AVN- 944, and brequinar from the ReFRAME library (**Table 3**) and valinomycin, rotenone, and mubritinib from the NPC library (**Table 4**), were the compounds with the strongest inhibitory activity. However, and similar to the results with VACV, the SI values of the compounds differed based on the CC_50_ values obtained from the MTT or XTT toxicity assays (Tables 3 and 4). These results demonstrated that a set of compounds from the ReFRAME and NPC libraries we previously identified as having antiviral activity against RNA viruses, also show potent antiviral activity against VACV and MPXV DNA viruses, expanding their potential as broad-spectrum antivirals against a diverse collection of RNA and DNA viruses.

**Figure 4.**
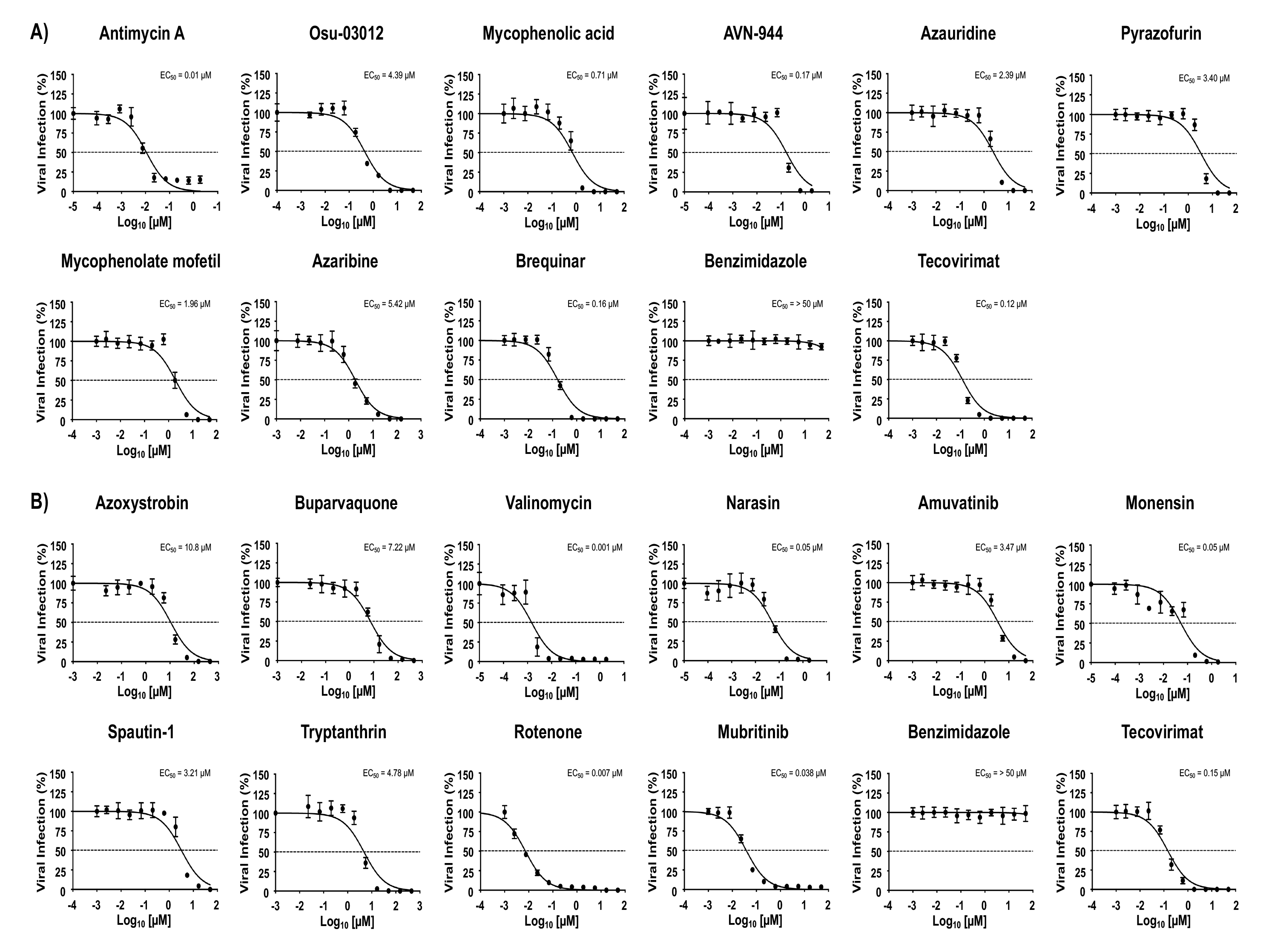
Antiviral activity of the ReFRAME and NPC compounds against MPXV: Human A549 cells (2×10^4^ cells/well, 96-well plates, quadruplicates) were infected with 200 PFU of MPXV and incubated with 3-fold serial dilutions (starting concentration of 50 µM) of the ReFRAME library compounds antimycin A, Osu-03012, mycophenolic acid, AVN-944, azauridine, pyrazofurin, mycophenolate mofetil, azaribine, or brequinar (**A**); or the NPC library compounds azoxystrobin, buparvaquone, valinomycin, narasin, amuvatinib, monensin, spautin-1, tryptanthrin, rotenone, or mubritinib (**B**). Benzimidazole and tecovirimat were included as negative and positive controls, respectively (**A** and **B**). Mock-infected cells and cells infected in the absence of drug were included as an internal control. Inhibition of viral replication was evaluated by quantifying the number of plaques at 24 hpi using a cross-reactive anti-VACV A33R polyclonal antibody and developed with an anti-rabbit Vectastain and DAB reagent. The EC_50_ of each compound was calculated using sigmoidal dose-response curves with GraphPad Prism. The dotted line indicates 50% inhibition. Data represent the means and SD from quadruplicates.

**Table 3.**
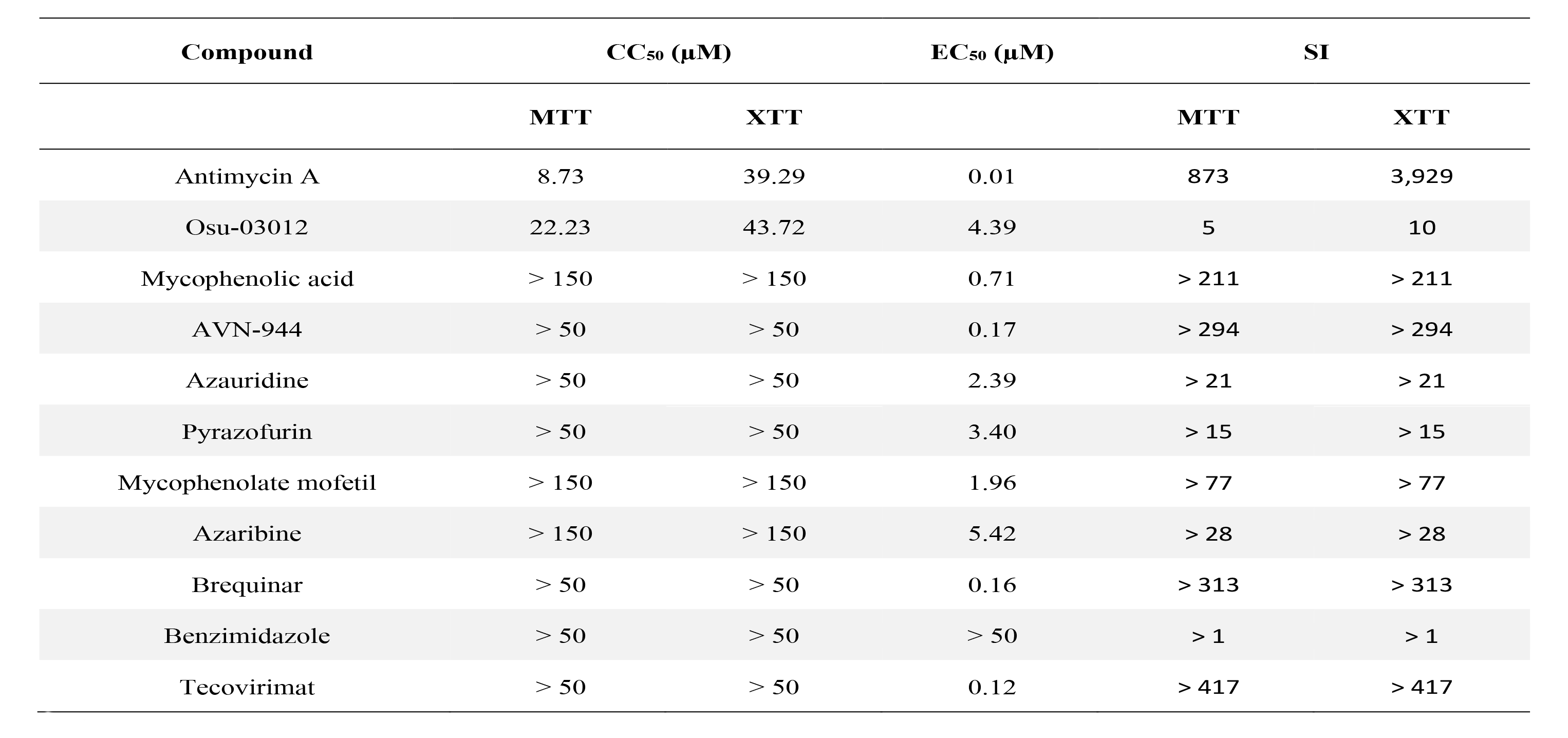
CC_50_, EC_50_, and SI of the ReFRAME antiviral compounds against MPXV.

**Table 4.** CC_50_, EC_50_, and SI of the NPC antiviral compounds against MPXV.

| Compound | CC<sub>50</sub> (μM) |  | EC<sub>50</sub> (μM) | SI |  |
| --- | --- | --- | --- | --- | --- |
|  | MTT | XTT |  | MTT | XTT |
| Azoxystrobin | 27.92 | > 450 | 10.8 | 3 | > 42 |
| Buparvaquone | 121.4 | > 450 | 7.22 | 17 | > 62 |
| Valinomycin | 0.29 | 13.63 | 0.001 | 290 | 13,630 |
| Narasin | 1.3 | 8.73 | 0.05 | 26 | 175 |
| Amuvatinib | 18.51 | 1.8 | 3.47 | 5 | 1 |
| Monensin | > 50 | > 50 | 0.05 | > 1,000 | > 1,000 |
| Spautin-1 | 8.00 | > 50 | 3.21 | 2 | > 16 |
| Tryptanthrin | 6.31 | 16.65 | 4.78 | 1 | 3 |
| Rotenone | 0.10 | > 50 | 0.007 | 14 | > 7,143 |
| Mubritinib | 0.07 | > 50 | 0.038 | 2 | > 1,316 |
| Benzimidazole | > 50 | > 50 | > 50 | > 1 | > 1 |
| Tecovirimat | > 50 | > 50 | 0.15 | > 333 | > 333 |

## DISCUSSION

The MPXV outbreak in early 2022 was declared by the WHO as a public health emergency of international concern, with over 30,286 confirmed cases and 38 deaths in the US as of March 2023 (86,746 cases and 112 deaths worldwide) (1-3). Alarmingly, the current MPXV outbreak exhibited uncommon person-to-person transmission through prolonged exposure or sexual contact with infected individuals (1) (5-7). Despite two vaccines being presently available for prevention of MPXV infection, insufficient supplies have restricted their accessibility to only those in high-risk populations, including immunocompromised individuals or men who have sex with other men. Only two antiviral drugs, tecovirimat and brincidofovir, previously approved by the US FDA for the treatment of smallpox, are available for the treatment of MPXV infection, highlighting the urgent medical need to identify novel antivirals for the treatment of MPXV and other Orthopoxvirus infections.

In this study, we first demonstrate the feasibility of using rVACV expressing both fluorescent (Scarlet and GFP) and luciferase (Nluc) proteins to easily identify compounds with antiviral activity against VACV. Next, we used these rVACV Nluc/Scarlet and rVACV Nluc-GFP to test the anti-VACV activity of compounds from the ReFRAME and NPC libraries we previously reported to have broad spectrum antiviral activity against RNA viruses (55-59). Using both fluorescent Scarlet or GFP and Nluc readouts we showed that several of the compounds in the ReFRAME and NPC libraries we tested had antiviral activity against VACV, and that assessing their antiviral activity using fluorescent (Scarlet or GFP) or luciferase (Nluc) expression resulted in similar EC_50_ and SI values.

Among the compounds in the ReFRAME library, antimycin A and AVN-944 were found to have the strongest antiviral activity against VACV. It is worth noting that antimycin A has been previously described to inhibit infection of multiple viruses, including porcine reproductive and respiratory syndrome virus (PRRRSV) (72), dengue virus (DENV) (73), IAV and IBV (64), equine encephalitis viruses (EEV), vesicular stomatitis virus (VSV), encephalomyocarditis virus (EMCV), Sendai virus (SeV), and hepatitis C virus (HCV) (74). AVN-944 has also been previously shown to have broad-spectrum antiviral activity against both RNA and DNA viruses, likely via reduction of GTP levels in infected cells (75-77). Notably, AVN-944 is currently in phase 1 human clinical trials for cancer treatment (78, 79). Other compounds with antiviral activity against VACV in the ReFRAME library included mycophenolic acid, pyrazofurin, mycophenolate mofetil, azaribine, and brequinar.

The NPC library compounds valinomycin, rotenone, and mubritinib also exhibited potent anti-VACV activity. Other compounds with antiviral activity against VACV from the NPC library included buparvaquone, narasin, and monensin. Valinomycin has broad-spectrum properties against tumors (80), bacteria (81), fungi (82), and viruses (83, 84), by acting as a potassium ionophore that increases potassium ion permeability of the mitochondrial inner membrane, thereby inhibiting oxidative phosphorylation (85). Monensin is also a selective ionophore that facilitates the transmembrane exchange of sodium ions for protons, which leads to the inhibition of intracellular transport of Golgi apparatus-associated proteins. Monensin has been shown to have anti-cancer (86) and antiviral properties (87-90), including VACV inhibition (91, 92). Rotenone has been extensively studied and validated as a complex I inhibitor of the mitochondrial electron transport chain (93). Rotenone toxicity in humans is moderate but can be uses therapeutically at doses posing very low safety risks, and has been studied as a potent broad-spectrum antiviral (94).

Importantly, six of the compounds identified in the ReFRAME library and all of the compounds from the NPC library to have antiviral activity against VACV were confirmed to also have antiviral activity against MPXV, supporting their potential use for the treatment of MPXV and potentially other Orthopoxvirus infections. Some of the differences observed in the compound’s antiviral activities against VACV and MPXV likely reflect different readout to assess antiviral activity. The ability of using fluorescence or luciferase reporter genes and the lower level of bio-contentment required to work with VACV support the use of the bireporter expressing rVACV Scarlet/Nluc and rVACV GFP/Nluc in HTS format to identify compounds with antiviral activity against Orthopoxviruses. Importantly, the compounds from the ReFRAME and NPC libraries we tested are FDA-approved for use in humans, which should facilitate the process to bring them into the clinic.

In summary, in this manuscript we have demonstrated that some compounds from the ReFRAME and NPC libraries previously identified to have antiviral activity against different RNA virus families, also have antiviral activity against VACV and MPXV, demonstrating their broad-spectrum antiviral potential for the treatment of different viral infections. Future studies in validated animal models of either VACV or MPXV (38, 71) will be necessary to assess the antiviral activity of the compounds *in vivo* and to test their therapeutic potential. It should be noted that the compounds we tested target host factors, which will pose a high genetic barrier to the emergence of viral escape mutants, a common event with antivirals directly targeting a viral protein activity. This is illustrated by tecovirimat resistance being the result of mutations in the viral gene encoding the target protein F13 (44). In addition, these results open the possibility of combination therapy using compounds targeting viral (e.g. tecovirimat) and host factors (e.g. ReFRAME and NPC compounds) for the treatment of Orthopoxvirus infections.

## ACKNOWLEDGMENTS

We want to thank BEI Resources for providing MPXV USA-2003 (NR-2500) and A33R polyclonal antibody (NR-628). This work was partly supported by grant PID2021- 128466OR-I00 funded by funded by MCIN/AEI/10.13039/501100011033 to M.L. and R.B.. Research in L.M-S laboratory is partially funded by the Antiviral Countermeasures Development Center, AC/DC (1U19AI171403-01); the Center for Antiviral Medicines & Pandemic Preparedness, CAMPP (1U19AI171443-01), and the QCRG Pandemic Response Program (1U19AI171110-01), three of the National Institute of Health (NIH) funded Antiviral Drug Discovery Centers for Pathogens of Pandemic Concern; by grants W81XWH2110103, W81XWH2110095, W81XWH1910496, and W81XWH2210283 from the Department of Defense (DoD) Peer Reviewed Medical Research Program (PRMRP); R43AI165089, R01AI161363, R01AI161175, R01AI145332, R01AI142985, R01AI141607, R21 AI173816, and 75N95022P00510 from the National Institute of Health (NIH); the Center for Research on Influenza Pathogenesis (CRIP), one of the National Institute of Allergy and Infectious Diseases (NIAID) funded Centers of Excellence for Influenza Research and Response (CEIRR; contract # 75N93021C00014); the a Systems Biology Lens (SYBIL; U19AI135972); the San Antonio Partnership for Precision Therapeutics, the San Antonio Medical Foundation, and the Texas Biomedical Research Institute Forum Foundation.

